# On the therapeutic potential of MAPK4 in triple-negative breast cancer

**DOI:** 10.1101/2022.08.24.505130

**Authors:** Fadia Boudghene-Stambouli, Mathilde Soulez, Natalia Ronkina, Anneke Dörrie, Alexey Kotlyarov, Ole-Morten Seternes, Matthias Gaestel, Sylvain Meloche

## Abstract

ERK3/MAPK6 (*MAPK6* gene) along with its paralog ERK4/MAPK4 (*MAPK4* gene) define a distinct subfamily of atypical mitogen-activated protein kinases (MAPKs)^1^. Much remains to be learned about the substrates and biological functions of these signaling enzymes. Interestingly, recent work has suggested that ERK4 promotes prostate cancer progression via the non-canonical activation of AKT/mTOR signaling^2,3^. In their recent study, Wang et al.^4^ report that ERK4 is expressed in a subset of triple-negative breast cancer (TNBC) cell lines and that this expression is critical for AKT activation and for sustaining TNBC cell proliferation *in vitro* and tumor growth in mice. They also show that depletion of ERK4 sensitizes TNBC cells to phosphatidylinositol-3-kinase (PI3K) inhibitors. They conclude that ERK4 is a promising therapeutic target for TNBC and has potential for combination therapy with PI3K inhibitors. Here, we raise concerns about the cellular models and experimental approaches used in this study, which compromises the conclusions on the oncogenic role of ERK4 in TNBC.

## RESULTS AND DISCUSSION

To investigate the role of ERK4 in TNBC, Wang et al.^4^ first surveyed a panel of TNBC cell lines and found that ERK4 is highly expressed in MDA-MB-231, Hs578T and HCC1937 cell lines (referred to as MAPK4-high), while other TNBC cell lines express low-to-undetectable levels of the kinase. They used the NSCLC cell line H1299 as a positive control of ERK4 expression. The MAPK4-high cell lines were then used throughout the study to validate the oncogenic role of ERK4. Compared to ERK3, which is expressed in most tissues and cell lines, ERK4 shows a more restricted expression profile and is generally found at low or undetectable levels in most cell lines. Previous independent observations from our laboratories suggested that ERK4 is not expressed in some of the MAPK4-high cell lines described in Wang et al.^4^ paper, prompting us to examine in more detail the expression of ERK4 in their cell line panel. We first measured the abundance of *MAPK4* mRNA by quantitative RT-PCR. We found that expression of *MAPK4* gene was undetectable in Hs578T and very low-to-undetectable (Ct > 35) in MDA-MB-231 and HCC1937 cells, but was reproducibly detected in HeLa and HEK 293T cells used as controls (Fig. 1a). No expression of *MAPK4* mRNA could be measured in the H1299 NSCLC cell line. In contrast, *MAPK6* expression was detected in all cell lines with Ct values ranging from 20 to 25 (Fig. 1a). Examination of internal transcriptomic data obtained by RNAseq analysis of MDA-MB-231 and Hs578T cells also revealed that *MAPK4* is not expressed to detectable levels in MDA-MB-231 and Hs578T cells, while *MAPK6* is expressed at significant levels with TPM values of 66.1 and 55.3, respectively (Fig. 1b). The DepMap portal uses a threshold of TPM ≥ 1 for calling an expressed gene (https://depmap.org/portal/). These data indicate that *MAPK4* gene is expressed at very low-to-undetectable levels in the MAPK4-high cell lines of Wang et al.^4^.

**Figure 1.**
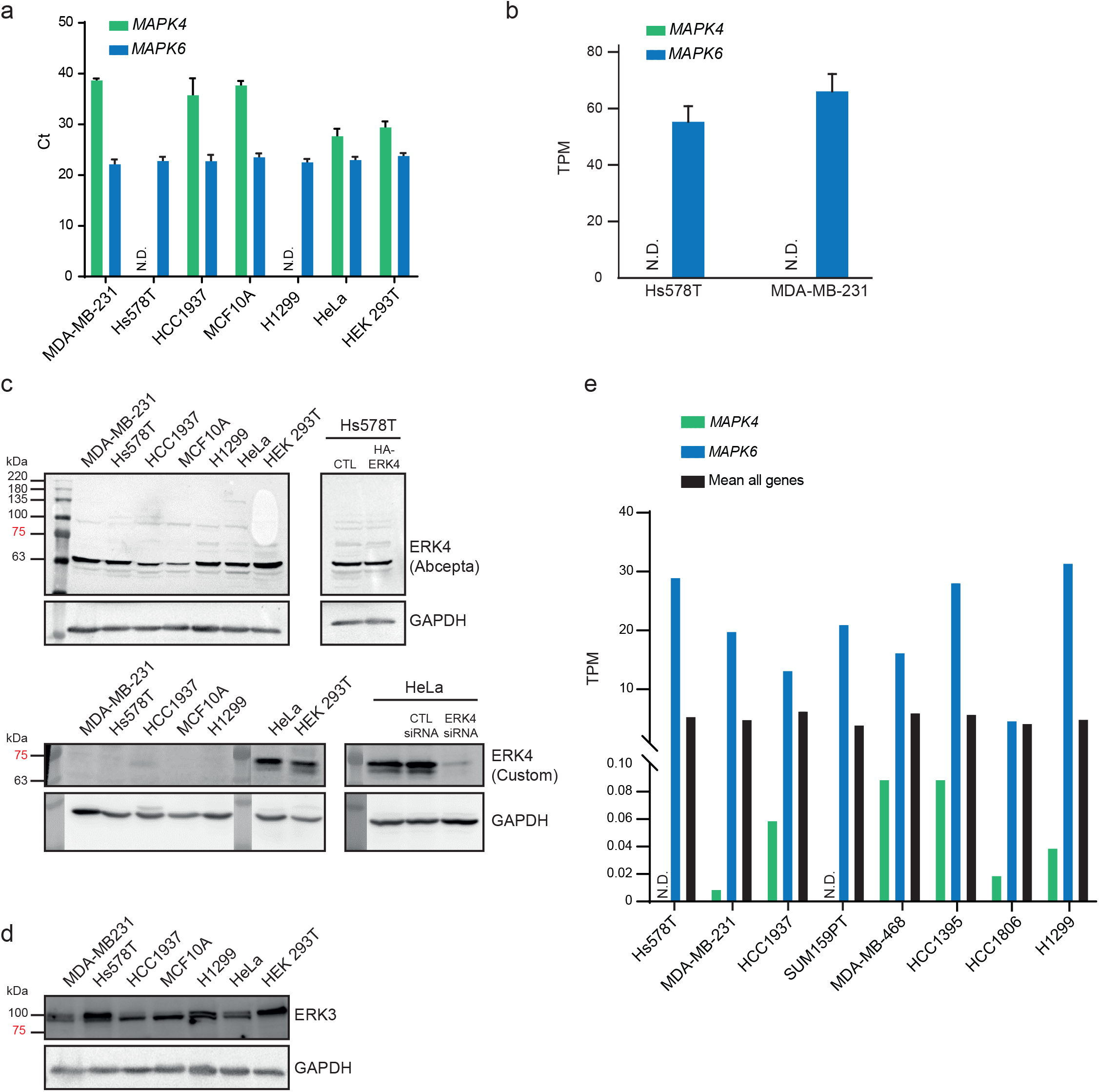
Expression of ERK4 mRNA and protein in TNBC cell lines. (**a**) Expression of *MAPK4* and *MAPK6* mRNA was measured by real-time qPCR in the human TNBC cell lines Hs578T, MDA-MB231 and HCC1937, the immortalized breast epithelial cell line MCF10A, and control H1299, HeLa and HEK 293T cells. Results are expressed as cycle threshold (Ct). Data are means ± SD (*n* = 3). (**b**) Expression of *MAPK4* and *MAPK6* mRNA in Hs578T and MDA-MB-231 measured by RNA-seq. Results are expressed as TPM and correspond to the mean ± SD of triplicate samples. **(c)** Expression of ERK4 and ERK3 protein was analyzed by western blotting using the indicated antibodies. To control for the specificity of the ERK4 antibodies, ERK4 was depleted from HeLa cells by siRNA, or ectopically expressed in Hs578T cells. GAPDH immunoblotting was used to control for protein loading. (**d**) Expression of ERK3 protein analyzed by western blotting. **(e)** Expression of *MAPK4* and *MAPK6* mRNA in human TNBC cell lines and H1299 NSCLC cell line extracted from the CCLE database. Results are expressed as TPM. Unprocessed original scans of blots are shown in Supplementary Fig. 1.

We next measured the expression of ERK4 protein in TNBC cell lines by western blotting. Two different antibodies were used for these experiments: the anti-MAPK4 from Abcepta (AP7298b) used in Wang et al.^4^ paper and a validated custom polyclonal ERK4 antibody that we use routinely in our laboratories. We failed to detect a specific ERK4 band in any of the cell lines, including Hs578T cells transfected with human ERK4 cDNA, using the anti-ERK4 antibody from Abcepta (Fig. 1c). Using our custom anti-ERK4 antibody, we observed an immunoreactive band of ∼70 kDa in control HeLa cells and HEK 293T cells (Fig. 1c). The signal intensity of the band was almost completely eliminated by siRNA-mediated depletion of ERK4 in HEK 293T cells. In agreement with the mRNA expression data, no expression of ERK4 protein was detected in MDA-MB-231, Hs578T and HCC1937 cell lines, or in control H1299 NSCLC cells (Fig. 1c). In contrast, ERK3 protein was expressed to detectable levels in all cell lines, in line with expression of the *MAPK6* gene (Fig. 1d).

The Cancer Cell Line Encyclopedia (CCLE) is a trusted database that compiles gene expression data for ∼1,000 cell lines available from public cell line repositories^5^. Interrogation of the CCLE database showed that *MAPK4* gene is expressed at very low or undetectable levels (TPM ≤1) in the TNBC cell lines MDA-MB-231, Hs578T, HCC1937, SUM159PT, MDA-MB-468, HCC1395 and HCC1806, and in the control NSCLC cell line H1299 (Fig. 1e). In contrast, *MAPK6* was expressed in all TNBC cell lines with TPM values ranging from 4.9 to 31.7. There was no correlation between the expression of *MAPK4* mRNA in CCLE TNBC cell lines and expression of ERK4 protein in TNBC cell lines of Wang et al.^4^. Taken together, these findings are inconsistent with a significant expression and oncogenic function of ERK4 in the TNBC cellular models used by Wang et al.^4^.

Another concern raised by the Wang et al.^4^ study relates to the proposed non-canonical activation of AKT by ERK4, which serves as rationale for the author’s hypothesis that ERK4 depletion will sensitize TNBC cells to PI3K inhibitors. ERK4 is a member of the MAPK family of Ser/Thr kinases, which are proline-directed kinases^6^. Accordingly, ERK4 phosphorylates its only known substrate MK5 on Thr 182, which is followed by a Pro residue^7,8^. The activation loop threonine of AKT1, AKT2 and AKT3 (Thr308 in AKT1) is not followed by a proline, raising questions about the validity and significance of this mechanism. In experiments performed prior to publication of the Wang et al.^4^ study, we have examined the ability of ERK4 to phosphorylate AKT on Thr308 and Ser473 in HEK 293T cells. We found that overexpression of either ERK4 or catalytically-inactive ERK4 KK49/50AA had no significant impact on the phosphorylation of AKT on its activating phosphorylation sites Thr308 and Ser473 (Fig. 2). In contrast, expression of ERK4, but not of catalytically-dead ERK4 KK49/50AA clearly induced the phosphorylation of MK5 on Thr182 and caused MK5-mediated phosphorylation of ERK4, which is evident by the retarded migration of ERK4 band^9^.

**Figure 2.**
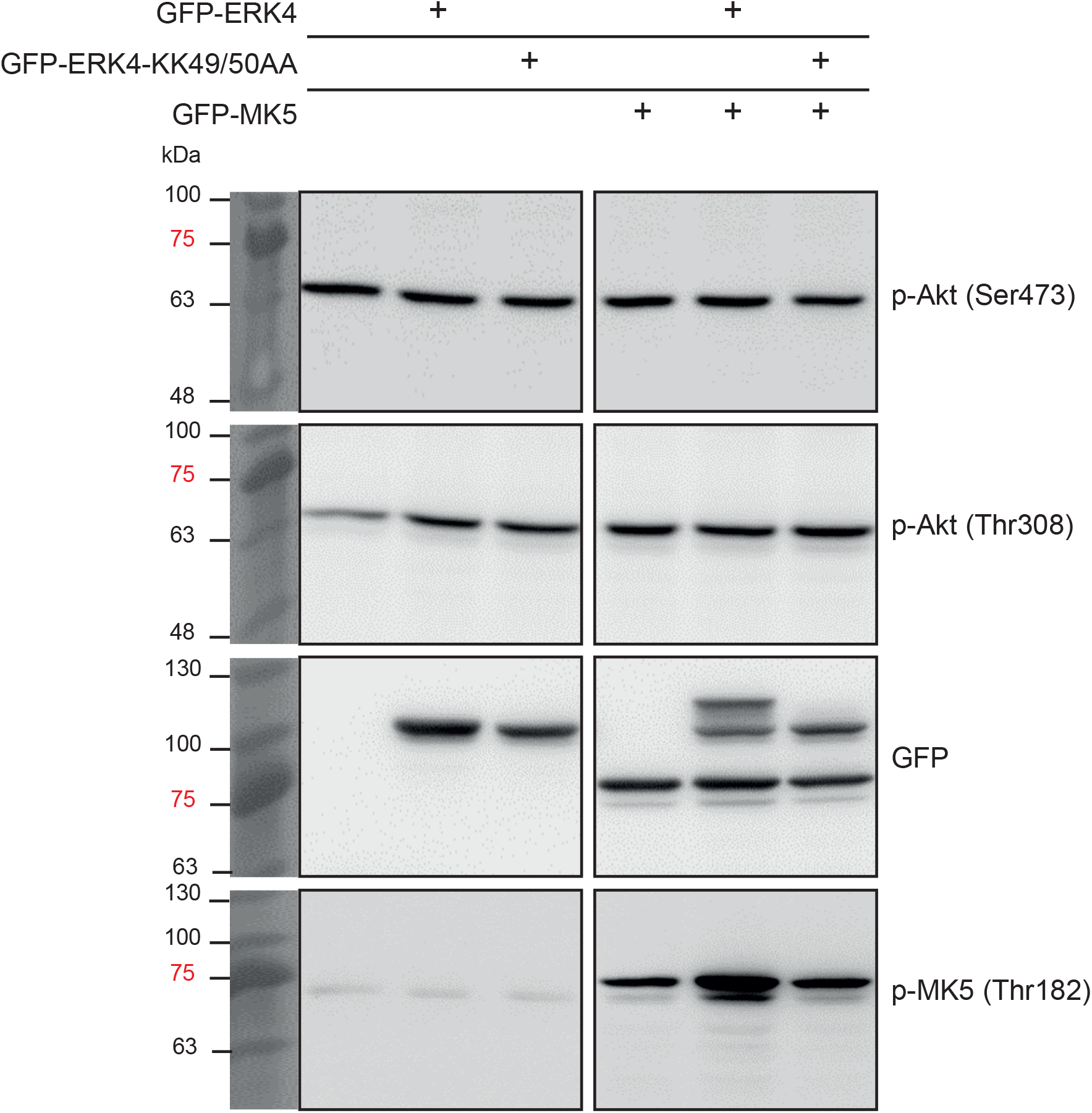
ERK4 catalytic activity has no significant impact on the phosphorylation of AKT on its activating phosphorylation sites Thr308 and Ser473. HEK 293T cells were transfected with GFP-ERK4 or catalytically-inactive ERK4 KK49/50AA with or without GFP-MK5. Phosphorylation of endogenous AKT on Thr308 and Ser473 or of ectopically expressed GFP-MK5 was analyzed by immunoblotting with phospho-specific antibodies. Expression of GFP-constructs was controlled with anti-GFP antibody. Unprocessed original scans of blots are shown in Supplementary Fig. 1.

In conclusion, our findings do not support the claim of Wang et al.^4^ that ERK4 promotes TNBC growth and is a promising therapeutic target for this cancer. We found no evidence that ERK4 is expressed to significant levels in the MAPK4-high cell lines used by the authors to validate the role of ERK4, thereby invalidating the conclusions of their study. In further support of a lack of role for ERK4, interrogation of the DepMap portal indicates that CRISPR/Cas9-mediated depletion of ERK4 in MDA-MB-232, Hs578T and HCC1937 has no effect on the proliferation of these cell lines, consistent with the lack of expression of the *MAPK4* gene (https://depmap.org/portal/gene/MAPK4?tab=overview). We also question the significance and generalization of the non-canonical activation of the AKT/mTOR pathway by ERK4 proposed by the authors. Our observations do not call into question the possible role of ERK4 in other solid cancers, which was not evaluated in this study.

## METHODS

### Cell lines and cell culture

All cell lines were obtained from the American Type Culture Collection. The TNBC cell lines MDA-MB-231, Hs578T and HCC1937 were further authenticated by short tandem repeat profiling (ATCC). MDA-MB-231, Hs578T and HEK 293T cells were cultured in DMEM supplemented with 10% fetal bovine serum (FBS) and antibiotics. H1299, HCC1937 and HeLa cells were cultured in RPMI supplemented with 10% FBS and antibiotics. MCF10A cells were cultured in DMEM/F12 supplemented with 10% FBS, epidermal growth factor, hydrocortisone, insulin and antibiotics. All cell lines were routinely tested for mycoplasma contamination.

### Transfections, infections, and RNA interference

For expression of GFP constructs, HEK 293T cells were transiently transfected with polyethylenimine and 1 µg of DNA per well in 12-well plates. After 24 h, the cells were lysed in the plate with 200 µl of 2X SDS-loading buffer per well. For siRNA treatment, 4×10^4^ HeLa cells were seeded into 12-well plates on the day before transfection. Next day, the cells were transfected with HiPerfect and 3 µl of 10 µM ERK4 siRNA from Santa Cruz Biotechnology (sc-62280) or control siRNA from Qiagen (1027280). After 48 h, the cells were lysed in the plate with 100 µl of 2X SDS-loading buffer.

For lentiviral infections, HEK 293T cells were transfected with polyethylenimine and 12 µg of plasmid DNA per 100-mm culture dish (3:2 ratio). For ectopic expression of ERK4, HEK 293T cells were transfected with packaging plasmids (pMD2/VSVG, pMDLg/pREE and pRSV/Rev) and pBabe-puro-HA-ERK4 or empty vector. After 48 h, virus-containing culture media was filtered and used to infect Hs578T cells. Polyclonal populations of infected cells were obtained by selection with puromycine at 2 µg/ml.

### Western blot analysis

Cell lysis and immunoblotting analysis were performed as described previously^10^. Custom polyclonal ERK4 antibody was raised in sheep against the full-length ERK4 protein produced in Sf9 cells and was affinity purified on CH-Sepharose covalently coupled to the protein antigen^7,11^. Commercial antibodies for immunoblotting were obtained from the following suppliers: anti-ERK4 (1/1,000; #AP7298b) from Abcepta; anti-ERK3 (1/1000; #ab53277) and anti-phospho-MK5(Thr182) (1/1,000; #ab138668) from Abcam; anti-GAPDH (1/2,000; #VMA00046), anti-HSC70 (1/2,000; #sc-7298), and anti-GFP (1/1,000; #sc-9996) from Santa Cruz Biotechnology; anti-phospho-AKT(Thr308) (1/1,000; #13038) and anti-phospho-AKT(Ser473) (1/1,000; #4051) from Cell Signaling Technology.

### Real-time quantitative PCR analysis

Total RNA was extracted with the RNeasy Mini Kit (Qiagen) and reverse transcribed with random primers and the High Capacity cDNA Archive Kit (Applied Biosystems) as described by the manufacturer. Quantitative real-time PCR was performed as previously described^12^. The following forward (F) and reverse (R) primers were used: *MAPK4* (F: tacggggagaatgctctttg; R: ccggattacagggatggtc), *MAPK6* (F: gtacaagttgatccccgaaaat; R: caaaagcaggatcctccaga), *HPRT* (F: tgatagatccattcctatgactgtaga; R: caagacattctttccagttaaagttg), and *GAPDH* (F: agccacatcgctcagacac; R: gcccaatacgaccaaatcc).

### Gene expression profiling

Total RNA was extracted from MDA-MB-231 and Hs578T cell lysates using RNAeasy purification kit (Qiagen). The quality of RNA was assessed on the Agilent 2100 Bioanalyzer and the RNA was quantified by QuBit fluorometry. Libraries (500 ng total RNA) were prepared using the KAPA Hyperprep messenger RNA (mRNA)-Seq Kit (KAPA Biosystems). Purified libraries were normalized by quantitative PCR (qPCR) using the KAPA Library Quantification Kit (KAPA Biosystems) and diluted to a final concentration of 10 nM. Sequencing was performed on the Illumina Nextseq 500 using the Nextseq500 High Output Kit (75 cycles) (read length 1 × 80 bp). Around 24–33 M single-end pass-filter reads was generated per sample. Library preparation and sequencing was done at the Institute for Research in Immunology and Cancer Genomics core facility. Alignment of RNA-seq reads was performed with the STAR aligner. The datasets generated during the current study are available from the corresponding author on reasonable request.

## ACKNOWLEDGMENTS

We thank R. Lambert for qPCR analysis and P. Gendron for help with bioinformatic analyses. S.M. is supported by grants from the Cancer Research Society and the Canadian Institutes for Health Research. M.G. is supported by Deutsche Forschungsgemeinschaft; O.-M.S. is supported by the Norwegian Cancer Society (grant 198119-2019) and Northern Norway Regional Health Authority/Helse Nord RHF (project HNF1547-20).

## AUTHOR CONTRIBUTIONS

F.B.-S., M.S., N.R., A.D., A.K. and O.-M.S. performed the experiments. M.G., O.-M.S. and S.M. conceived and supervised the studies. S.M. wrote the manuscript. All authors commented on the manuscript.

