## Supplementary figures and images for "On the therapeutic potential of MAPK4 in triple-negative breast cancer"

### Supplemental Figure 1

#### Supplementary Figure 1

1c

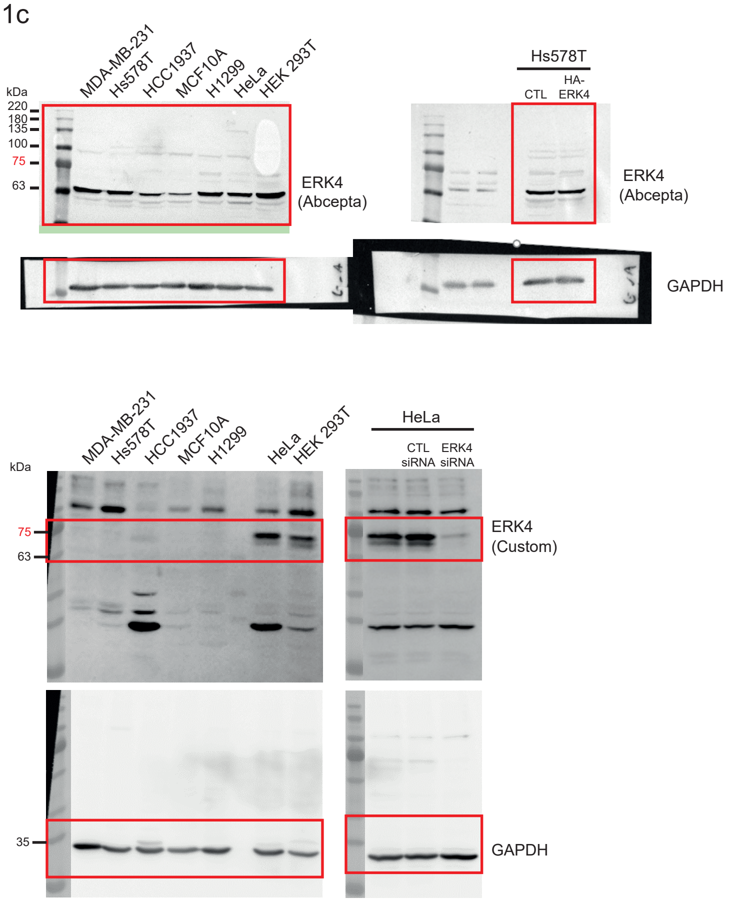


1d

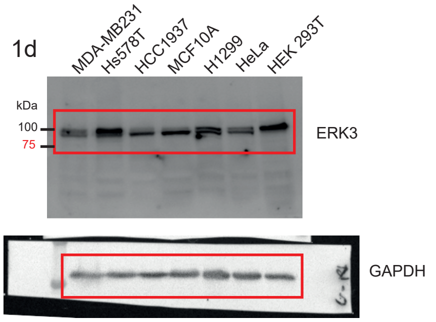


2

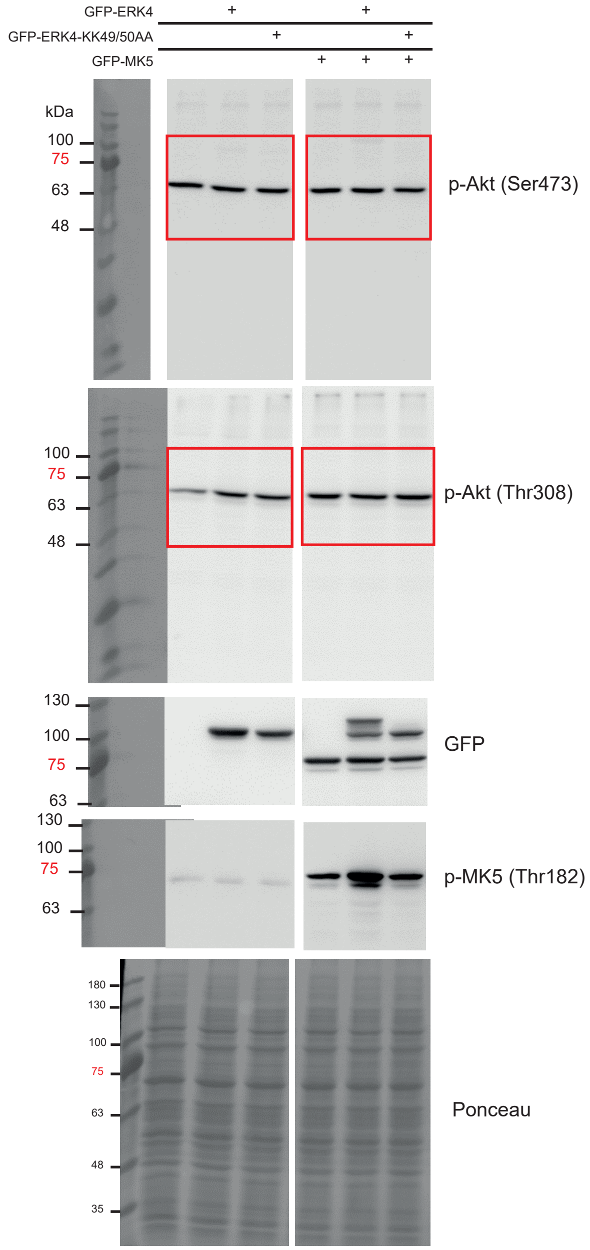
